# Independent human mesenchymal stromal cell-derived extracellular vesicle preparations differentially affect symptoms in an advanced murine Graft-versus-Host-Disease model

**DOI:** 10.1101/2020.12.21.423658

**Authors:** Rabea J. Madel, Verena Börger, Robin Dittrich, Michel Bremer, Tobias Tertel, Nhi Ngo Thi Phuong, Hideo A. Baba, Lambros Kordelas, Jan Buer, Peter A. Horn, Astrid M. Westendorf, Sven Brandau, Carsten J. Kirschning, Bernd Giebel

**Affiliations:** Institute of Medical Microbiology, University Hospital Essen, University of Duisburg-Essen, Essen, Germany; Department of Infectious Diseases, University Hospital Essen, University of Duisburg-Essen, Essen, Germany; Institute of Transfusion Medicine, University Hospital Essen, University of Duisburg-Essen, Essen, Germany; Institute of Pathology, University Hospital Essen, University of Duisburg-Essen, Essen, Germany; Department of Hematology and Stem Cell Transplantation, University Hospital Essen, University of Duisburg-Essen, Essen, Germany; Department of Otorhinolaryngology, University Hospital Essen, University of Duisburg-Essen, Essen, Germany

**Keywords:** mesenchymal stem cells, mesenchymal stromal cells, extracellular vesicles, exosomes, Graft versus Host Disease

## Abstract

Extracellular vesicles (EVs) harvested from cell culture supernatants of human mesenchymal stromal cells (MSCs) suppress acute inflammation in preclinical models of various diseases. Furthermore, they promote regeneration of damaged tissues. Following successful clinical treatment of a steroid-refractory Graft-versus-Host-Disease (GvHD) patient with EVs prepared from conditioned media of human bone marrow (BM)-derived MSCs, we aim to improve MSC-EV production and quality control towards clinical application. Observing functional differences of independent MSC-EV preparations *in vitro*, we established an optimized murine GvHD model for the analysis of independent MSC-EV preparations *in vivo*. To this end, T cell depleted allogeneic BM cells co-transplanted with naïve allogeneic spleen-derived T cells induced GvHD symptoms with reproducible strengths in mice being preconditioned by ionizing irradiation. Administration of MSC-EV preparations with confirmed *in vitro* immune modulatory properties at three consecutive days significantly suppressed GvHD symptoms. In contrast, application of MSC-EV preparations lacking these *in vitro* immune modulating capabilities failed to suppress GvHD symptoms. Thus, our results reveal therapeutic differences among independent MSC-EV preparations that had been produced in a standardized manner. Thus, given this functional heterogeneity, any individual MSC-EV preparation considered for the clinical application should be evaluated for its potency prior to administration to patients.

## Introduction

Allogeneic hematopoietic stem cell transplantation (alloSCT) is the most frequently applied cell-based therapy. For now, it is conducted to treat more than 70 malignant and non-malignant hematologic diseases, including leukaemia and lymphoma (Sweeney and Vyas, 2019). In many cases, it is considered as the only therapy offering curative perspectives. However, alloSCT remains associated with severe side effects. Approximately 35% of recipients receiving matched related donor transplants and up to 50% of those receiving unrelated or alternative donor transplants develop mild (grade I and II) to severe manifestations (grade III and IV) of Graft-versus-Host Disease (GvHD). In GvHD, donor immune cells attack and destroy previously healthy tissues of the recipient. Severe acute GvHD (aGvHD) is associated with a high mortality, with only 25-40% of patients with grade III aGvHD and 1-2% of patients with grade IV aGvHD surviving more than two years (Malard et al., 2020).

First-line treatment to prevent GvHD involves permanent immunosuppression including systemic anti-inflammatory compound, mainly corticosteroid (methylprednisolone) administration. However, such anti-GvHD treatments often fail and cause severe side effects. Approximately, 50% of acute or chronic GvHD patients react sufficiently while the others are steroid refractory. Notably, patients with steroid refractory acute GvHD have a dismal long-term prognosis with survival rates of only 5-30% (Zeiser and Blazar, 2017). There is no accepted second-line treatment for steroid refractory GvHD. This is because most studies, which analysed the efficacy of other compounds as GvHD therapy, are retrospective, single-armed or have produced inconsistent results (Malard et al., 2020). Thus, in view of the constantly growing number of alloSCTs (Passweg et al., 2020), the heavy burden of the disease and the lack of effective anti-GvHD therapies novel treatment strategies for steroid-refractory GvHD are urgently needed.

At the turn of the millennium, mesenchymal stem/stromal cells (MSCs) had been shown to mediate immunosuppression (Bartholomew et al., 2002; Di Nicola et al., 2002). Subsequently, Le Blanc and colleagues successfully explored the capability of BM-derived MSCs to suppress GvHD symptoms in a severe treatment resistant grade IV acute GvHD patient (Le Blanc et al., 2004). Consequently, numerous centres started to apply MSCs for steroid-refractory aGvHD treatment with varying outcomes (Baron and Storb, 2012). Although several groups confirmed the beneficial effect of MSCs application to acute GvHD patients in a proportion of patients, a phase III clinical trial failed to show efficacy (Galipeau, 2013). Apparently, not all administered MSCs have the capability to suppress aGvHD symptoms successfully.

In contrast to the initial hypothesis that MSCs act in cellular manners, a collection of observations implied, MSCs mediate their preclinical and clinical effects rather in paracrine modes (Caplan and Dennis, 2006). Indeed, in 2009 and 2010, EVs were shown to execute the MSCs’ function in an acute kidney injury or a myocardial infarction model, respectively (Bruno et al., 2009; Lai et al., 2010). Subsequently, we started to investigate the MSC-EVs’ immunomodulatory capabilities and confirmed their ability to modulate pro-inflammatory immune responses *in vitro*. Considering their therapeutic potency and having had a treatment-refractory acute GvHD patient whose immune cell functions were successfully modulated by MSC-EVs *in vitro*, an individual MSC-EV treatment scheme for this patient was developed; allogeneic MSC-EVs were applied in seven escalating doses over a period of two weeks (Kordelas et al., 2014). Remarkably, following treatment, GvHD symptoms were massively suppressed for more than 4 months (Kordelas et al., 2014). In order to consecutively translate MSC-EVs into regular GvHD clinic, we progressively improved our MSC-EV production platform. However, working with primary BM-derived MSCs provides several challenges. MSCs are limited in their expansion capabilities; consequently, EVs need to be prepared from conditioned media of MSCs of varying donors. Since MSCs are a very heterogeneous cell entity (Phinney, 2012; Phinney et al., 1999; Radtke et al., 2016; Vogel et al., 2003), we have compared the pro- and anti-inflammatory content of resulting EV preparations right from the beginning and observed huge differences among independent MSC-EV preparations. For example, significant differences within cytokine profiles of four independent, but identically produced MSC-EV preparations were recorded (Kordelas et al., 2014).

To avoid failures of the MSC field as exemplified by a failed phase III clinical trial of MSC treated GvHD patients (Galipeau, 2013), we consider functional testing of any MSC-EV preparation as essential prior to clinical application. Currently, the knowledge of the concrete mode(s) of action of MSC-EV preparations is sparse, challenging the establishment of appropriate *in vitro* assays for the potency prediction of independent MSC-EV preparations. To identify MSC-EV preparations being potent to suppress GvHD symptoms, we thus established and applied an advanced murine GvHD model. Subsequently, we compared the capability of independently produced human MSC-EV preparations to modulate the GvHD symptoms and observed correlations with results obtained in a novel type of a mixed lymphocyte reaction (MLR) assay; MSC-EV preparations that reduced the content of activated CD4^+^ T cells in the MLR assay also reduced the disease severity in GvHD mice. In contrast, MSC-EVs lacking recognizable impacts on CD4^+^ cells within the MLR assay also failed to improve the GvHD symptoms in GvHD mice. Thus, the adapted and improved GvHD model provides important information about the potency of individual MSC-EV preparations and helps to qualify *in vitro* assays for the future potency testing of given MSC-EV preparations.

## Materials and methods

### MSC growth and expansion

Human BM aspirates from healthy donors were obtained following informed consent according to the Declaration of Helsinki. Their usage was approved by the ethics committee of the University of Duisburg-Essen (12-5295-BO). To raise MSCs, aliquots of obtained BM aspirates were seeded into cell culture flasks containing endothelial basal media (EBM-2, Lonza, Cologne, Germany) supplemented with 10% human platelet lysate (PL; produced in house) and the provided bullet kit (includes human endothelial growth factor ([EGF], hydrocortisone, gentamicin, amphotericin-b [GA-1000], vascular endothelial growth factor [VEGF], human fibroblast growth factor [hFGF], insulin-like growth factor [R3-IGF-1], ascorbic acid and heparin). After incubation for 24 hours at 37°C in a 5% CO_2_ atmosphere, non-adherent cells were removed by medium exchange to DMEM low glucose (PAN Biotech), supplemented with 10% PL, 100 U/mL penicillin-streptomycin-glutamine (Thermo Fisher Scientific, Darmstadt, Germany) and 5 IU/mL Heparin (Ratiopharm, Ulm, Germany). Cultures were continuously cultured at 37°C in a 5% CO_2_ atmosphere and regularly screened microscopically until the first MSC colonies became visible. Following trypsin/ EDTA (Sigma-Aldrich, Taufkirchen, Germany) treatment including a washing step, adherent cells were re-seeded at densities of approximately 1000 cells/cm into 4-layer stack cell factory™ systems (Thermo Fischer Scientific). Within the second passage, MSCs were analysed according to the criteria of the *International Society of Cell and Gene Therapy* (ISCT) (Dominici et al., 2006). Briefly as described before, MSCs were fluorescently labelled with anti-CD14, anti-CD31, anti-CD34, anti-CD44, anti-CD45, anti-CD73, anti-CD90, anti-CD105 and anti-HLA-ABC antibodies (suppl. Table 1) and analysed by flow cytometry (Cytoflex; Software Cytexpert 2.3, Beckman-Coulter, Krefeld, Germany). Upon passage 3, the MSCs’ osteogenic and adipogenic differentiation potentials were confirmed by conventional MSC differentiation assays (Kordelas et al., 2014; Radtke et al., 2016). Upon reaching densities of approximately 50% confluency, conditioned media (CM) were changed every 48 hours. At 80% confluency, MSCs were passaged. For the preservation of the CM, cells and larger debris were removed by 2000 × g centrifugation of cell suspensions for 15 minutes (Rotor: JS-5.3; Beckman Coulter). MSC-free CM were stored at −20°C until usage. CMs were screened regularly for mycoplasma contamination (Venor®GeM OneStep, Minerva Biolabs, Berlin, Germany).

**Table 1:**
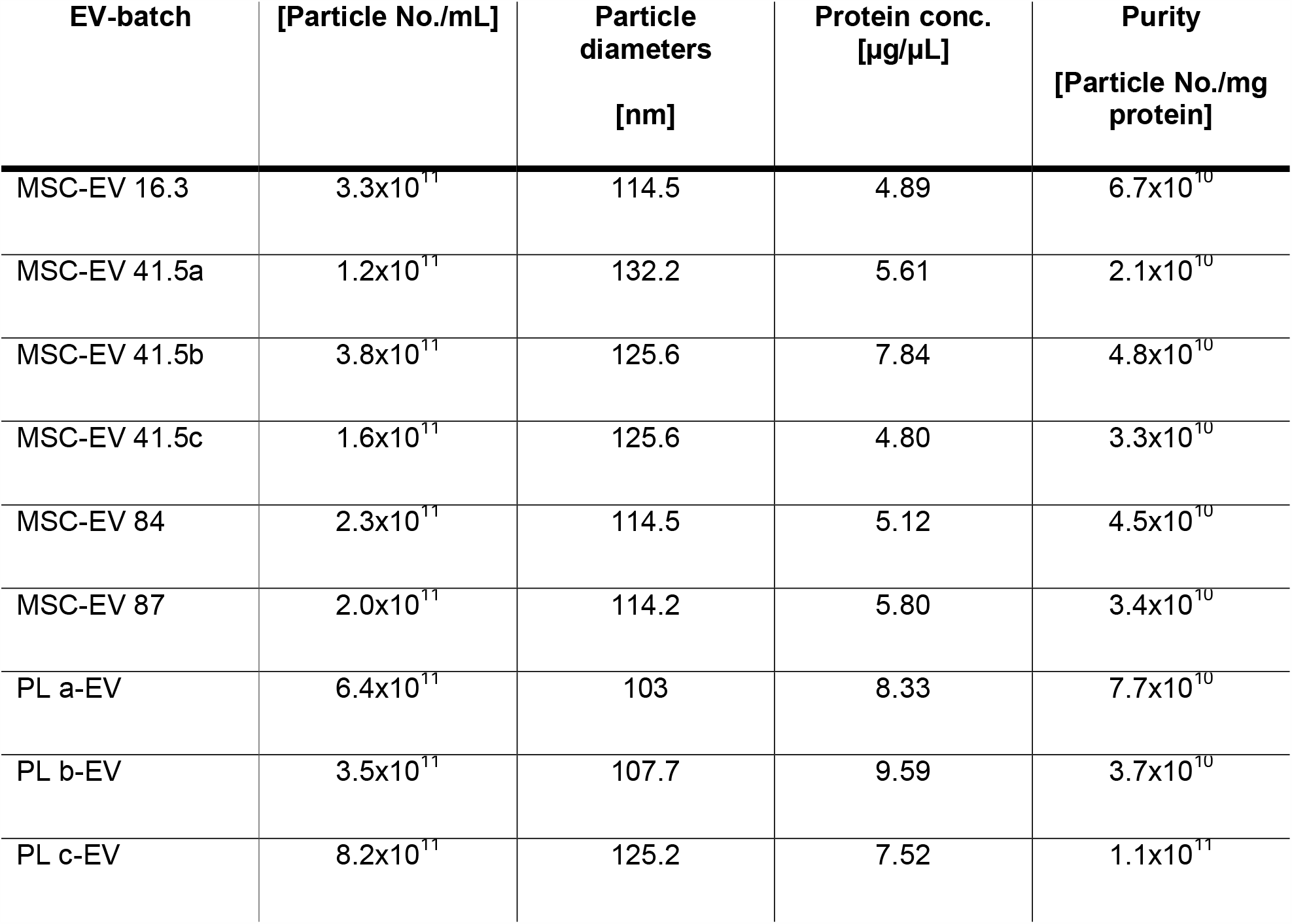
EV preparation characteristics: Particle No./mL and particle diameters (nm) as measured by NTA, Protein concentration as measured by BCA and the calculated purity index, particle No. per 1 mg protein.

| EV-batch | [Particle No./mL] | Particle diameters<br>[nm] | Protein conc.<br>[µg/µL] | Purity<br>[Particle No./mg protein] |
| --- | --- | --- | --- | --- |
| MSC-EV 16.3 | $3.3 \times 10^{11}$ | 114.5 | 4.89 | $6.7 \times 10^{10}$ |
| MSC-EV 41.5a | $1.2 \times 10^{11}$ | 132.2 | 5.61 | $2.1 \times 10^{10}$ |
| MSC-EV 41.5b | $3.8 \times 10^{11}$ | 125.6 | 7.84 | $4.8 \times 10^{10}$ |
| MSC-EV 41.5c | $1.6 \times 10^{11}$ | 125.6 | 4.80 | $3.3 \times 10^{10}$ |
| MSC-EV 84 | $2.3 \times 10^{11}$ | 114.5 | 5.12 | $4.5 \times 10^{10}$ |
| MSC-EV 87 | $2.0 \times 10^{11}$ | 114.2 | 5.80 | $3.4 \times 10^{10}$ |
| PL a-EV | $6.4 \times 10^{11}$ | 103 | 8.33 | $7.7 \times 10^{10}$ |
| PL b-EV | $3.5 \times 10^{11}$ | 107.7 | 9.59 | $3.7 \times 10^{10}$ |
| PL c-EV | $8.2 \times 10^{11}$ | 125.2 | 7.52 | $1.1 \times 10^{11}$ |

### Preparation of EVs

For EV harvesting, CMs were thawed and further purified following 45 min 6,800 × g centrifugation (Rotor: JS-5.3) by a subsequent 0.22 µm filtration step using rapid flow filter (Nalgene, Thermo Fisher Scientific). EVs were precipitated in 10% polyethylene glycol 6000 (PEG) and 75 mM sodium chloride (NaCl) by overnight incubation and subsequent centrifugation at 1,500 × g and 4°C for 30 min as described previously (Kordelas et al., 2014; Ludwig et al., 2018). Pelleted EVs were re-suspended and washed with sterile 0.9% NaCl solution (Braun, Nelsungen, Germany) to remove contaminating soluble proteins. Next, EVs were re-precipitated by ultracentrifugation at 110,000 × g for 130 min (XPN-80, Ti45 rotor, k-factor: 133). Finally, EV pellets were re-suspended in 10 mM HEPES 0.9% NaCl buffer (Thermo Fisher Scientific). Concentration was adjusted so that 1 mL final sample contained the EV yield prepared from CM of approximately 4 × 10^7^ MSC equivalents. MSC-EV preparations were stored at −80°C. Repetitive thawing and freezing cycles were avoided. For control purposes, fresh PL supplemented media were processed in parallel (including incubation for 48 hours at 37°C, 5% CO_2_, saturated water vapour atmosphere).

### Physical and protein-biochemical analyses of MSC-EV preparations

MSC-EV preparations were characterized according to the *minimal information for studies of extracellular vesicles* 2018 (MISEV2018) commitment (Thery et al., 2018). The particle concentration and their average sizes within obtained MSC-EV preparations were determined by nanoparticle tracking analysis (NTA) on a ZetaView^®^ platform (ParticleMetrix, Meerbusch, Germany) as described before (Ludwig et al., 2018; Sokolova et al., 2011). The device was calibrated using a polystyrene bead standard (100 nm, Thermo Fisher Scientific). Samples were loaded and videos were recorded at all 11 positions, with 5 repetitions. Additional settings included sensitivity 75, shutter 75, minimum brightness 20, minimum size 5, and maximum size 200. The median value (X50) for size was used for data analysis.

The protein contents of the MSC-EV preparations were determined using the bicinchoninic acid (BCA) assay (Pierce, Rockford, IL, USA) in 96-well plates according to the manufacturer’s recommendations.

### Western Blotting

Western blot analyses were performed as described before (Ludwig et al., 2018). The following antibodies were used: anti-syntenin (clone EPR8102; Abcam, Cambridge, U.K.), anti-prohibitin (clone II-14-10; Thermo Fisher Scientific), anti-calnexin (ab10286; Abcam), anti-CD81 (clone JS-81; BD Biosciences, San Jose, CA, U.S.A.), anti-CD9 (clone VJ1/20.3.1; kindly provided by Francisco Sánchez, Madrid, Spain) and anti-CD63 (H5C6; BioLegend, San Diego, CA, USA). Band intensities were analysed using Image J.

### Transmission electron microscopy

Transmission electron microscopy (TEM) analyses were performed representatively and had been described and published before (Wang et al., 2020).

### Mixed lymphocyte reaction assay

To test allogeneic immune responses, a novel multi-donor mixed lymphocyte reaction assay (MLR) was used (Bremer et al., in preparation). Briefly, peripheral blood mononuclear cells (PBMC) of 12 donors were harvested from buffy coats via conventional Ficoll density gradient centrifugation (Beckmann et al., 2007; Giebel et al., 2004), pooled, and stored in the vapour phase of liquid nitrogen. Upon thawing, PBMCs were cultured in RPMI 1640 medium (Thermo Fisher Scientific) supplemented with 10% human AB serum (produced in house) and 100 U/mL penicillin and 100 µg/mL streptomycin (Thermo Fisher Scientific). Mixed PBMCs were plated at densities of 600,000 cells per 200 µL and per well of 96-well u-bottom shape plates (Corning, Kaiserslautern, Germany) and cultured either in the presence or absence of MSC-EV preparations to be tested at 37°C in a 5% CO_2_ atmosphere. After 5 days, cells were harvested, stained with a collection of specifically selected fluorescent labelled antibodies (CD4-BV785 [300554, Clone: RPA-T4, BioLegend]), CD25-PE [12-0259-42, Clone: BC-96, Thermo Fischer Scientific] and CD54-AF700 [A7-429-T100, Clone: 1H4, EXBIO]) and analysed on a Cytoflex flow cytometer (Software Cytexpert 2.3, Beckman-Coulter). Activated and non-activated CD4^+^ T cells were discriminated by means of their CD25 and CD54 expression, respectively. 25 µg of MSC-EV preparations to be tested were applied into respective wells.

### Mouse breeding and experimentation

Inbred C57Bl/6 strain mice of specific genotypes were bred in house (Kruger et al., 2015). Wild-type female Balb/c mice were purchased from Jackson Laboratories (Bar Harbour, ME, USA) or Charles River Laboratories (Sulzfeld, Germany). Specifically, 12 to 14 weeks old Balb/c mice were used as recipients and C57Bl/6 mice as grafts of bone marrow to be purified and transplanted. All mice were housed in a specific pathogen-free facility and had access to autoclaved food and drinking water ad libitum. 7d prior to preconditioning ionizing irradiation (IIR), drinking water and food pellets were mixed/soaked with antibiotics (neomycin, ampicillin, vancomycin, and metronidazole, each at 0.33 g/l) and provided until termination of the experiments. All animal procedures were performed in accordance with the international guidelines for good laboratory practice and approved by the North Rhine-Westphalia State Agency for Nature, Environment and Consumer Protection (LANUV, reference numbers 84-02.04.2011.A319 and 84-02.04.2014.A494).

### Bone marrow T cell depletion

BM cells of C57Bl/6 donor mice were collected by flushing tibias and femurs by using 10 mL syringes (Terumo) filled with cell culture medium (DMEM, 10% FCS, 1% Penicillin-Streptomycin, 1% Antibiotic-Antimycotic, Thermo Fisher Scientific) (Kruger et al., 2015). After red blood cell lysis by incubation in lysis buffer (Roth) 1:10 for 3 min at RT, Thy1.2 cells were depleted via negative selection using the CD90.2 MicroBeads mouse Kit (Miltenyi Biotec, Bergisch Gladbach, Germany) according to the manufacturer’s instructions.

### Naïve T cell purification from spleens

For isolation of naïve CD4^+^ T cells, spleens from C57Bl/6 mice were smashed through a 70 µm cell strainer (Thermo Fisher Scientific). After red blood cell lysis (1:10 dilution for 3 min at RT, Roth), naïve CD4^+^ T cells were purified via negative selection using the Naïve CD4^+^ T cell Isolation mouse Kit (Miltenyi Biotec) according to the manufacturer’s recommendations.

### Experimental GvHD model

Recipient Balb/c mice were fed with antibiotic containing drinking water and antibiotic soaked chow starting five days (d5) prior to preconditioning. To induce GvHD at d0, recipient mice total bodies were subjected to a minimally lethal dose of 8 Gy of X-rays (source RS 320, X-Strahl Ltd., operated at 300 kV, 10 mA at a dose rate of 161.55 cGy/min). On d1 recipient mice received intravenously 5 × 10^6^ T cell depleted C57Bl/6 BM cells and 0.7 × 10^6^ purified C57Bl/6 splenic naïve CD4^+^ T cells. The symptoms of the GvHD mice were treated with aliquots of selected MSC-EV preparations at 3 consecutive days, namely at d7, d8, and d9, applied i.v. as 300 µL sterile saline suspensions. Information about the dosing are provided in the result section.

Mouse weights and clinical scores were documented daily until principal experiment termination at d11. Towards clinical scoring, five parameters were monitored (Cooke et al., 1996), namely weight change, posture (hunching), activity, fur texture, and skin integrity wherein each value’s maximal score was 2. Animals reaching the cumulatively maximal score of 10 were sacrificed immediately while otherwise scarification was performed by enforced isofluran inhalation at d11. Blood was sampled immediately by retro-orbital bleeding and collected in both, EDTA-tubes (VWR) for blood cell counting and Heparin-tubes (BD) to gain plasma for cyto- and chemokine content analysis. Specifically, bead-based luminex assay systems (R&D Systems, Minneapolis, USA) specific to KC, TNF, IL-6, and G-CSF were applied according to the manufacturer’s protocols on a Luminex 200 instrument equipped with the XPonent software (Luminex, Austin, USA).

### Colon analyses

Mice were sacrificed via isoflurane inhalation on day 11 and the colons were prepared and flushed with PBS to remove faeces. The colon was carefully flattened with a thin wooden stick, rolled up into a Swiss Role as described before (Moolenbeek and Ruitenberg, 1981) and fixed in 4% formalin in a tissue cassette. The samples were then embedded in paraffin and 5 μm thin sections were cut via microtome. These tissue sections were then stained with hematoxylin and eosin and analysed by light microscopy. The pathology was evaluated and graded from 0-4, with 0: no pathology and 4: severe pathology, based on crypt and microvilli integrity (histological score).

Regulatory T cells within colons were investigated as reported previously (Pastille et al., 2019). Briefly, prepared colons were flushed with PBS, cut into pieces and washed with PBS containing 3 mM EDTA (Sigma-Aldrich) for 10 min at 37°C. Thereafter, two washing steps with RPMI-1640 (Gibco) containing 1% FCS, 1 mM EGTA and 1.5CmM MgCl_2_ (Sigma-Aldrich) were performed. Subsequently, the colon pieces were digested in RPMI-1640 medium containing 20% FCS and 100 U/mL collagenase IV (Sigma-Aldrich). Single cells were separated from the remaining tissue by filtration through 40 µm cell strainers and washed with RPMI-1640 thereafter. Cells were surface-stained with Fixable Viability Dye (FVD) eFluor 780 (Thermo Fisher Scientific) and anti-CD4-PE antibodies (clone H129.19, BD Biosciences). Afterwards, intracellular staining with anti-FoxP3-FITC antibodies (clone: FJK-16s; Thermo Fischer Scientific) was performed with the BD Cytofix/Cytoperm Kit (BD Biosciences) according to the manufacture instructions. Labelled cells were analysed on a FACS Canto flow cytometer (BD Biosciences). Tregs were identified as FVD^-^CD4^+^FoxP3^+^ cells.

### Statistical analysis

Prism 8 (GraphPad Software, San Diego, USA) was used for data presentation and statistical analysis. Differences between groups were determined by one-way analysis of variance (one-way ANOVA) with Tukey’s, Dunnett’s or Kruskal-Wallis multiple comparisons test, comparing the mean of each group with the mean of every other group. P-values of less than 0.05 were considered to represent statistical significance.

## Results

### Co-transplantation of allogeneic T cell-depleted BM and splenic naïve T cells robustly induces GvHD pathology

Intending to optimize the production of MSC-EVs for the treatment of steroid refractory acute GvHD patients, we set up a murine GvHD model for the preclinical testing of obtained MSC-EV preparations. To mimic the allogeneic bone marrow transplantation setting and to become able to control the severity of the subsequent GvHD responses, we chose a strategy combining the stratifying aspects of previously described protocols, i.e. the co-transplantation of T cell depleted BM and a defined number of isolated naïve T cells (Anderson et al., 2003; Riesner et al., 2016).

C57Bl/6:H-2Kb mice were chosen as recipients and T cell depleted bone marrow cells from Balb/c:H-2Kd mice as the basal component of the allogeneic transplants. In order to trigger experimental GvHD by allogeneic BM transplantation in a controlled manner, distinct numbers of naïve CD4^+^ T cells freshly purified from spleens of C57Bl/6 graft mice were co-transplanted with 5 × 10^6^ T cell depleted C57Bl/6 BM cells into recipient Balb/c mice that were myeloablated by total body IIR 24 h before BM transplantation. Five different groups were investigated that either were co-transplanted with none, 0.125, 0.25, 0.5 or 1.0 × 10^6^ CD4^+^ cells. For controlling the irradiation effectiveness, additional mice were irradiated without receiving any transplants. Expectedly, their weight loss was that high that they had to be sacrificed preterm at d9 following irradiation (Figure 1A, B).

**Figure 1:**
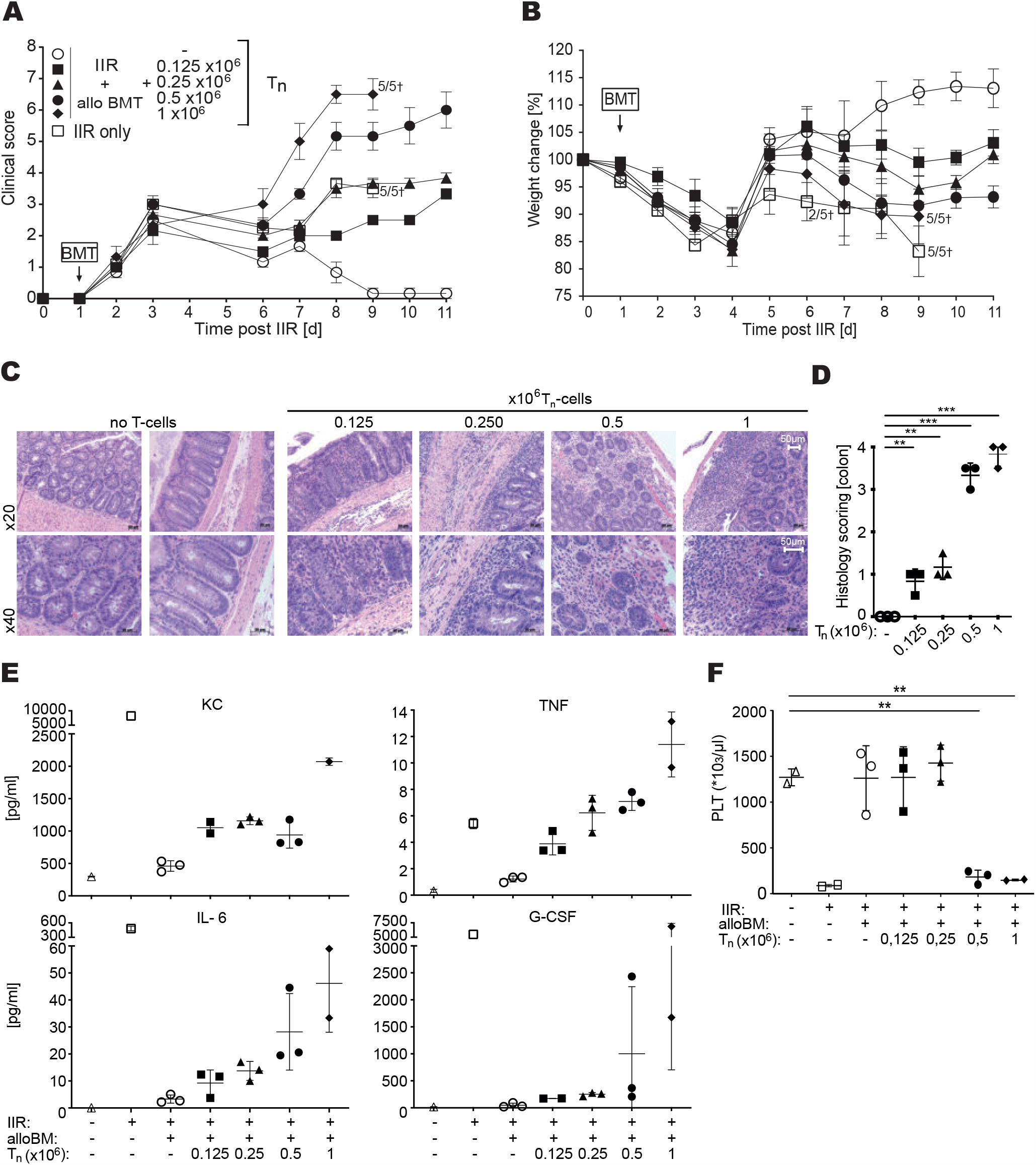
Experimental murine GvHD severity depends on the amount of allogenic T cells co-transplanted with T cell depleted BM cells. (A) clinical score and (B) percentage weight change of experimental groups (n=5 animals per group) that received indicated numbers of splenic CD4^+^ T cells. The irradiation effect was controlled on irradiated mice not receiving allogeneic BM transplants (IIR only). (C) haematoxilyn and eosin stained colon sections of a representative animal of each transplantation group. (D) histology score based on the assessment of the haematoxilyn and eosin stained colon sections of the mice of the different groups at the day of sacrification (n=3). (E) Plasma cytokine concentration and (F) platelet count in whole blood (n=3 per group). Tn, cotransplanted T cell numbers; alloBM, allogeneic T cell-depleted bone marrow cells; IIR, ionizing irradiation; BMT, bone marrow cell transplantation; PLT, platelets; mean values ± SEM; ANOVA ***p<0,001; **p<0,01; *p<0,05).

Following transplantation, mice were scored every 24 h; specifically, body weight, habitus, activity, fur and skin appearance were monitored. As expected, the number of T cells co-transplanted with the transplant correlated with the severity of GvHD symptoms (Figure 1A). While mice transplanted with BM only recovered completely from a temporal weight reduction until d11, GvHD severity, as indicated by weight loss, increased with the number of co-transplanted T cells (Figure 1B). Host animals co-transplanted with 1 × 10^6^ T cells developed such strong GvHD symptoms that they had to be sacrificed prior to the scheduled experimental endpoint at d11 (Figure 1A, B).

In line with the previous result, the number of co-transplanted T cells correlated with the severity of colon GvHD symptomatology. Specifically, loss of crypts and microvilli as well as the numbers of apoptotic cells and intestinal wall swelling increased with the T cell dose; mice which received 1 × 10^6^ CD4^+^ cells for example virtually lacked all crypts and microvilli (Figure 1C, D). Blood analyses revealed the same tendency, the number of co-transplanted T cells correlated with the serum content of the pro-inflammatory cytokines keratinocytes-derived chemokine (KC, CXCL1), tumour-necrosis factor alpha (TNFα), interleukin-6 (IL-6) and granulocyte colony stimulating factor (G-CSF) (Figure 1E) and inversely correlated with the number of platelets per blood volume (Figure 1F). Overall, the obtained results qualify the allogeneic co-transplantation of T cell depleted BM and defined numbers of purified splenic naïve T cells as a tuneable strategy to robustly and reproducibly induce GvHD symptoms in mice.

### Applied MSC-EV preparations can modulate the severity of the GvHD symptoms in the established aGvHD model

To test whether MSC-EV preparations can modulate the severity of GvHD symptoms in the established mouse model, we decided to co-transplant 5 × 10^6^ T cell depleted C57Bl/6 BM cells together with 0.7 × 10^6^ pure naïve T cells. For the *proof-of-principle* experiments, we chose a MSC-EV preparation (MSC-EVs 41.5a) that had revealed therapeutic potency in a murine ischemic stroke model before (Wang et al., 2020). At day 7, 8 and 9 post transplantation mice were either treated with aliquots of the MSC-EV 41.5a preparation being equivalent to the EV-harvest from the supernatant of 1.2 × 10^6^ MSCs, or with PBS. Mice were continuously monitored until sacrification at day 11 post transplantation (Figure 2A). The phenotypic appearances of animals were documented (Figure 2B). Blood samples as well as gut biopsies were analysed (Figure 2C, D). In contrast to PBS treated aGvHD mice, MSC-EV 41.5a treated aGvHD mice increased their weight and had an overall healthier appearance (Figure 2B). Furthermore, the platelet number in the blood of MSC-EV treated mice was increased in comparison to PBS treated aGvHD mice and the morphologies of their guts were massively improved (Figure 2C-E). Thus, the aGvHD symptoms of the mice can successfully be modulated by an appropriate MSC-EV preparation.

**Figure 2:**
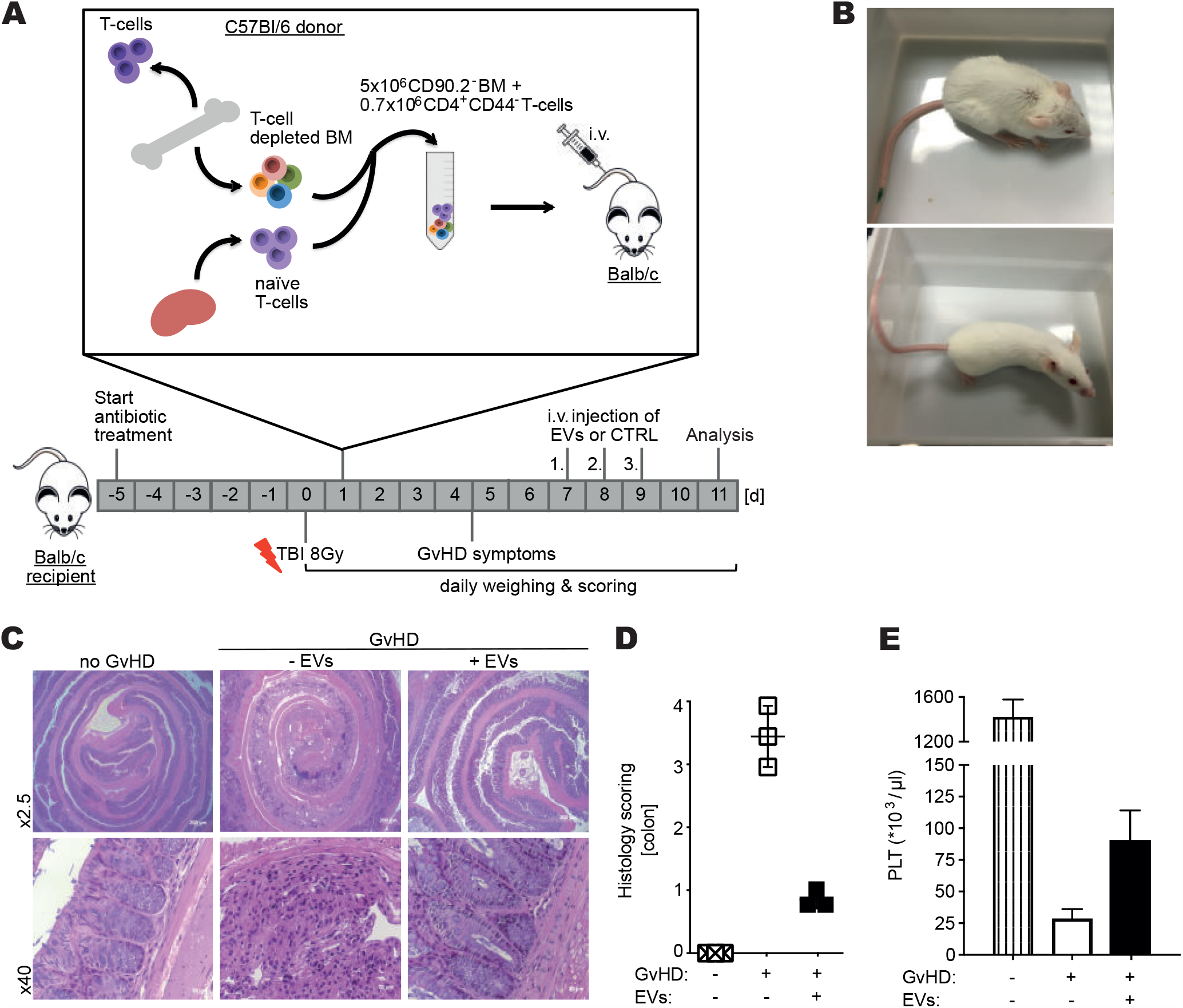
MSC-EV application can reduce experimentally induced murine GvHD symptoms. (A) Experimental design: Recipient Balb/c mice were preconditioned by total body irradiation (TBI) with 8 Gy. On d1 post irradiation, 5 × 10^6^ Thy1.2^-^ BM cells were co-transplanted with 0.7 × 10^6^ splenic naïve CD4^+^ T cells (CD4^+^CD44^-^ T cells), both harvested from C57Bl/6 mice. At consecutive d7, d8 and d9 the MSC-EV 41.5a preparation or PBS as control were administered i.v. (EV yield of conditioned media of 1.2 × 10^6^ MSCs was diluted in PBS to final injection volumes of 300 µL). Mice were monitored and individually scored for GvHD symptoms daily and sacrificed on d11. (B) Representative comparison of the habitus and fur integrity of a PBS treated GvHD mouse (upper panel) and a MSC-EV 41.5a preparation treated GvHD mouse (lower panel) at d11. (C) representative haematoxilyn and eosin stained colon sections of healthy control and PBS (-EVs) or MSC-EV 41.5a preparation treated (+EVs) GvHD mice. (D) histological score based on the assessment of the haematoxilyn and eosin stained colon sections and (E) peripheral blood platelet (PLT) counts of healthy control and PBS or MSC-EV 41.5 preparation treated GvHD mice (n=3). Mean values ± SD.

### Independent MSC-EV preparations differ in their in vitro immunomodulatory capabilities

aGvHD is largely allogeneic immune response driven. To mimic this aspect of aGvHD pathology *in vitro*, based on the concept of a previous publication (Pachler et al., 2017), we have set up a modified multi-donor human leukocyte mixed lymphocyte reaction (MLR) assay that can be used to reproducibly evaluate the immunomodulatory potential of individual MSC-EV preparations. Briefly, peripheral blood mononuclear cells (PBMCs) prepared from buffy coats of healthy peripheral blood donors (n=12) are pooled, aliquoted, and cryopreserved until usage. After thawing, 0.6 × 10^6^ pooled cells are seeded to individual wells of 96 well plates and cultured for 5 days in the presence or absence of MSC-EV preparations to be tested. Due to the mutual allogeneic stimulation within the PBMC mixture reproducibly high portions of CD4^+^ T cells get activated. Phenotypically, the activated CD4^+^ T cells are recognized as CD25^+^CD54^+^ cells. Here, we used this novel MLR assay to analyse whether capabilities of selected MSC-EV preparations to modulate allogeneic T cell responses *in vitro* associate with their potency to suppress GvHD symptoms *in vivo*.

In total, we prepared EVs from conditioned media of MSCs obtained from four different donors (MSC 16.3, 41.5, 84 and 87). All MSCs fulfilled the *bona fide* ISCT criteria (suppl. Figure 1) (Dominici et al., 2006). To study potential batch-to-batch variations, we also prepared EVs from conditioned media of MSCs 41.5 which were raised in three times independently from the same donor MSC batch (MSC-EVs 41.5a, 41.5b, 41.5c). Particle concentrations of the MSC-EV preparations as measured by NTA varied from 1.2 × 10^11^ to 3.8 × 10^11^ particles per mL MSC-EV preparation, which corresponded to the particle yield of supernatants of 4 × 10^7^ MSCs (Table 1). The protein concentration varied from 4.80 to 8.38 mg/mL resulting in particle concentrations per mg protein ranging from 2.1 × 10^10^ to 6.7 × 10^10^ (Table 1). Thus, the particle and protein concentrations of obtained MSC-EV preparations are all in a similar range. The average size distribution of prepared particles ranged from 107.9 to 132.2 nm (Table 1). All MSC-EV preparations contained the tetraspanins CD9, CD63 and CD81 and according to the expectations lacked Calnexin (Suppl. Figure 2).

Although none of the parameters appeared to be ideal for the normalization of sample amounts to be applied into the MLR assays, in favour of good tradability we decided to apply volumes of the selected MSC-EV preparations containing 25 µg proteins. As MSCs are raised in human platelet lysate supplemented media, which had not been EV depleted, EV preparations of fresh PL supplemented media (PL-EVs) served as controls (Table 1).

In the absence of any EV preparation, MLR assay cultures on average contained 37.6% (31.3-40.3%) activated CD4^+^ T cells (Figure 3). The application of 25 µg of the PL-EVs control sample did not significantly alter the amount of activated T cells (Figure 3). Also, no severe impacts on the percentage of activated T cells were recorded when MSC-EV preparations 41.5b (36.5%), 16.3 (32.3%) or 84 (37.4%) were applied (Figure 3). In contrast, a clear reduction in activated T cell contents was observed when MSC-EV preparations 41.5a (12.2%), 41.5c (21.3 %) or 87 (15.0%) were applied (Figure 3). Thus, results of the MLR analysis demonstrate, independent MSC-EV preparations, even if they derive from independent batches of the same donor material, can considerably differ in their *in vitro* immunomodulatory capabilities.

**Figure 3:**
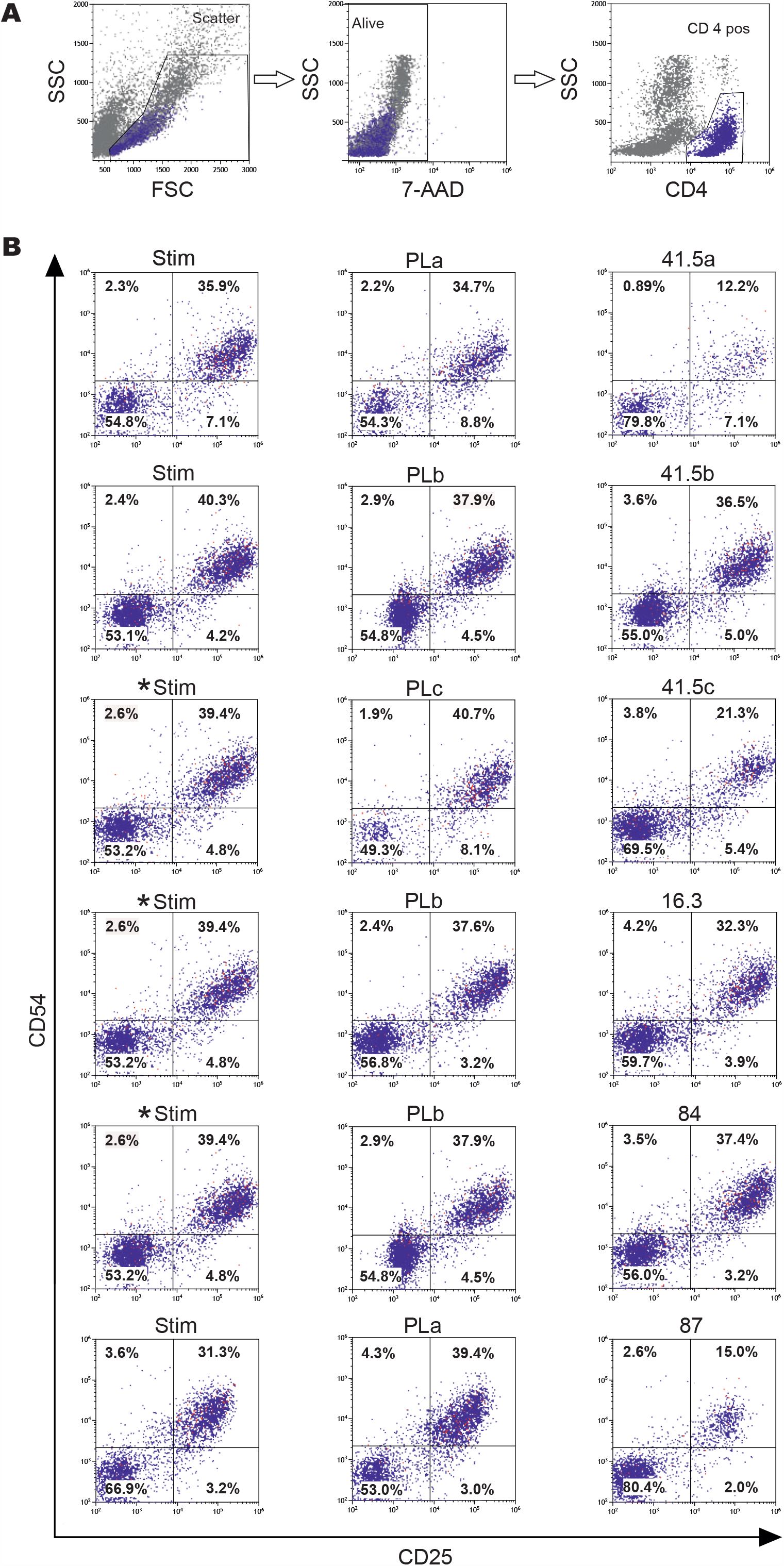
Independent MSC-EV preparations differ in their *in vitro* immunomodulatory capabilities. EV preparations either obtained from human platelet lysate supplemented non-conditioned (PLa-c) or MSC-conditioned media (41.5a-c, 16.3, 84, 87) were tested for their immunomodulatory capabilities in a novel multi-donor MLR assay. Specifically, approx. 600.000 cells of a mononuclear cell mixture derived from peripheral blood samples of 12 healthy donors were cultured for 5 days, either in the presence or absence of 25 µg of given EV preparations. Thereafter, cells were harvested and labelled with antibody cocktails containing anti-CD4, anti-CD25 and anti-CD54 antibodies. The proportions of activated CD4^+^ T cells (CD25^+^CD54^+^) were determined by flow cytometric analyses. CD4^+^ cells were gated according to their light scatter features and as 7AAD^-^, CD4^+^SSC^low^ cells (blue, first row). *: controls are identical (respective EV preparations were analyses in parallel at the same day).

### MSC-EV preparations with in vitro immunomodulatory capabilities modulate GvHD symptoms in vivo

Next, we evaluated whether the *in vitro* immunomodulatory properties of the given MSC-EV preparations reflect their therapeutic functions to suppress aGvHD symptoms in the optimized GvHD mouse model. Aliquots of five different MSC-EV preparations (MSC-EV preparations 16.3, 41.5a, 41.5b, 41.5c and 87) and two PL-EV preparations (PLb and PLc) were applied according to the established experimental aGvHD protocol. Each individual EV preparation was applied to 5 to 10 mice with established aGvHD symptoms. In addition, aliquots of all *in vivo* applied samples were analysed in parallel on Western blots either being sequentially probed with anti-Syntenin and anti-Calenxin, or anti-CD9, anti-CD81 and anti-CD63 antibodies, respectively (suppl. Figure 2).

The results of the *in vivo* studies revealed that the MSC-EV preparations 41.5a, 41.5c and 87 improved the aGvHD clinical and gut histological scores significantly (Figure 2A-E). In contrast, administration of MSC-EV 41.5b preparation was largely ineffective in modulating aGvHD symptoms. For unknown reasons MSC-EV preparations 16.3 increased the aGvHD symptoms in 4 out of 10 animals that, as a consequence thereof, had to be sacrificed prematurely (Figure 4A, B). Intestinal analyses had been representatively performed for mice, which were treated with the MSC-EV preparations 41.5c and 16.3, in the latter case only mice were analysed that were not prematurely sacrificed. Following administration of the MSC-EV 41.5c preparation, the colon crypts of 3 out of 3 mice had almost the same appearance as non-GvHD control mice. In contrast, administration of MSC-EV 16.3 preparation had no significant effect on the gut morphology in 3 out of 3 mice; as in the PBS treated control animals the colon crypts were almost completely destroyed and the guts revealed a huge proportion of fibrotic tissue (Figure 4D, E).

**Figure 4:**
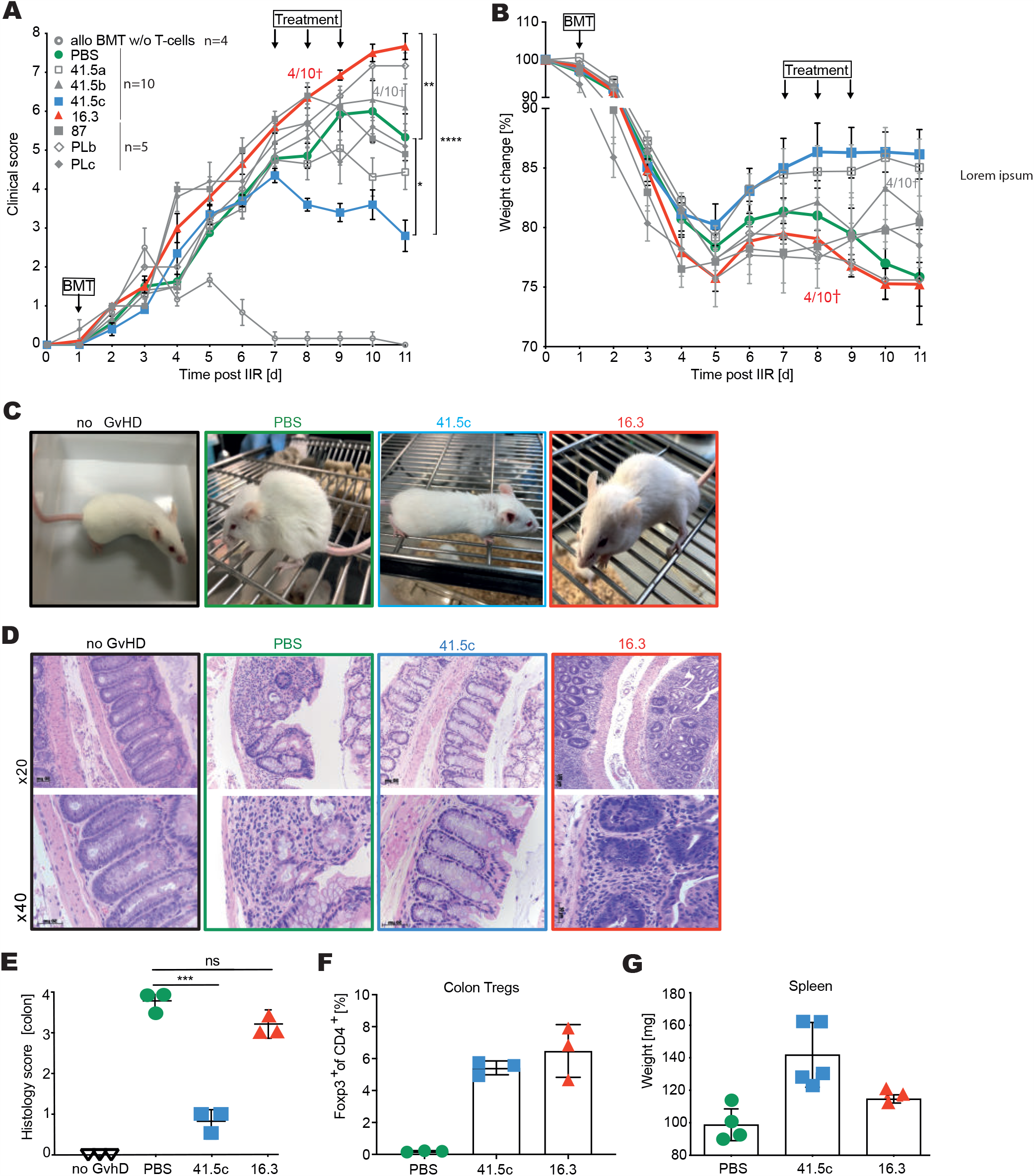
Independent MSC-EV preparations differ in their ability to inhibit GvHD symptoms *in vivo*. (A) Comparative analysis of mice treated with 8 independent MSC-EV preparations; blue, representative for a therapeutic effective MSC-EV preparation; red, for a therapeutic ineffective MSC-EV preparation; green, PBS control. Clinical scoring based on 5 criteria [weight change, posture, activity, fur texture, and skin integrity], the weight is separately illustrated (B). (C-G) Comparison of therapeutic effects of MSC-EV 41.5c and 16.3 preparations on aGvHD symptoms: (C) Representative posture of a normal (no GvHD) and GvHD-diseased mice either treated with (PBS) or MSC-EVs. (D) representative haematoxilyn and eosin stained colon sections of healthy control (no GvHD) and of PBS and MSC-EV preparation treated GvHD mice. (E) histological score based on the assessment of the haematoxilyn and eosin stained colon sections; (F) proportion of regulatory T cells (Tregs) among the population of colon residing CD4^+^ T cells and (G) spleen weight of PBS and MSC-EV treated GvHD mice. (E-H) represents mean +/-SD of n = 3 mice per group. The data are representative for one out of 3 independent experiments. BMT, bone marrow and T cell transplantation. ns, not significant. ANOVA ***p<0,001, ** p<0,01, *p<0,05.

Next, to test whether the improvement of the gut morphology following application of MSC-EV 41.5c preparation is accompanied by the presence of regulatory T cells, the frequency of Foxp3^+^ CD4^+^ T cells (Treg) was determined (Figure 4F). Indeed, following MSC-EV application more Tregs were recovered in the intestines of 6 out of 6 GvHD mice, remarkably irrespectively of whether the MSC-EV preparations revealed activities in the MLR assay (Figure 3) or whether the overall aGvHD symptoms were improved. We also determined the weight of spleens of MSC-EV treated and control GvHD mice. Following treatment with the MSC-EV 41.5c preparation mice had up to 4 times higher spleen weights than GvHD mice treated with the MSC-EV 16.3 preparation (Figure 4G). Thus, although MSCs were raised under identical cell culture conditions and EVs prepared with the same protocol, our obtained data confirm functional differences among independent MSC-EV preparations as well as functional batch-to batch variations.

### MSC-EV preparations modulate GvHD symptoms in dose dependent manners

So far, all MSC-EV preparations were applied at doses of 1.2 × 10^6^ cell equivalents per mouse. With an average body weight of 25 g this reflects the EV equivalents of ∼4.5 × 10^7^ cells per kg body weight. In contrast to the doses that were applied here, we previously applied roughly 5 × 10^5^ cell equivalents per kg body weight of the treated patient (Kordelas et al., 2014). Thus, compared to the successfully treated aGvHD patient 100 fold increased MSC-EV doses were applied to the aGvHD mice. To evaluate dose effects of the applied MSC-EV preparations, the active MSC-EV 41.5c and the inactive MSC-EV 16.3 preparations were applied in comparison to the selected dose in 3-fold lower or 3 fold-higher doses, respectively.

While, reduction of the MSC-EV 41.5c preparation dose reduced the strength of the observed therapeutic effect, the application of the 3 time doses did not reveal any further improvement (Figure 5A). This implies that threshold activities exist which had been reached for the MSC-EV 41.5c preparation with the original dose. As expected, reduction of MSC-EV 16.3 preparation had no impact on the GvHD symptoms, while the 3 time dose slightly improved the symptoms almost to the same extent as the 1/3 dose of MSC-EV 41.5c preparation (Figure 5 A, B).

**Figure 5:**
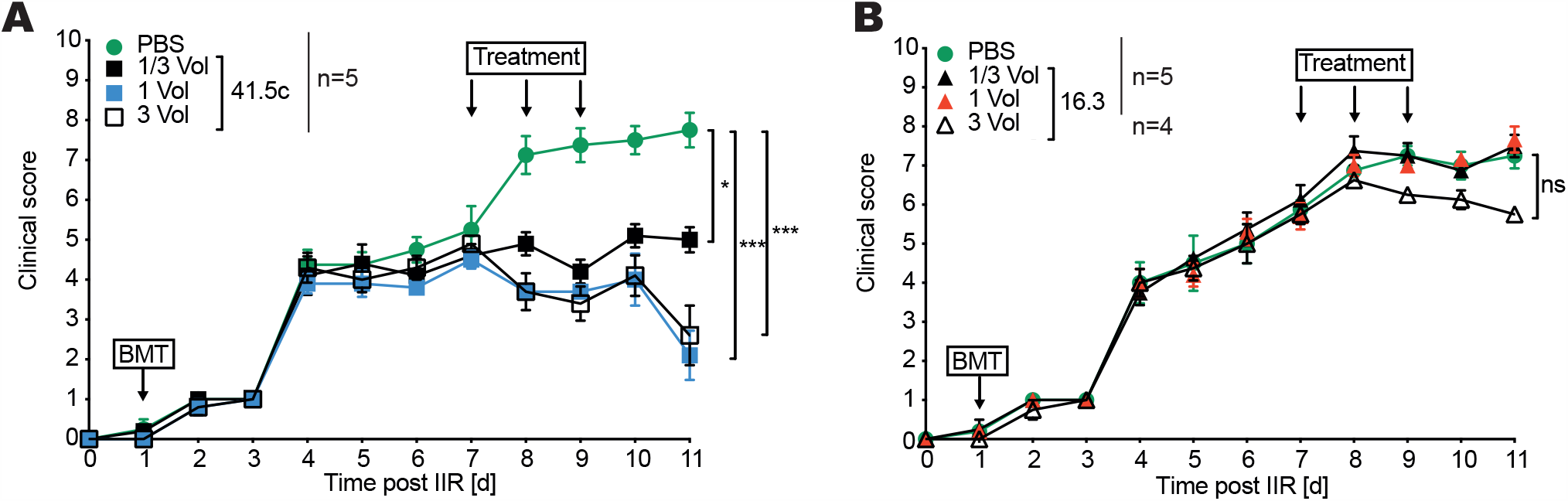
MSC-EV preparations modulate GvHD symptoms in dose dependent manners. (A, B) Clinical score upon triple, single and third dose administration of (A) MSC-EV 41.5c and (B) 16.3 preparations to experimental aGvHD diseased mice according to the regimen represented in Figure 2A. Mean values +/- SD of n=8-10. The graphs depict results of one out of two independent experiments with comparable outcomes. IIR, ionizing irradiation; ns, not significant; ANOVA ***p<0,001, ** p<0,01, *p<0,05).

In speculatively extrapolating these results, we would assign an at least 9 fold higher therapeutic activity to the MSC-EV 41.5c preparation than to the MSC-EV 16.3 preparation. Attempting to identify a potential surrogate marker, we compared several EV-specific parameters, including particle and protein concentration and the intensities of the specific CD9, CD63, CD81, syntenin and calnexin bands in Western blots, but could not detect any coincidence (Table 1; suppl. Table 2; suppl. Figure 2): The MSC-EV 41.5c preparation revealed a comparable protein concentration than the MSC-EV 16.3 preparation but showed a lower particle concentration (Figure 6A). In contrast, the intensities of the bands of the exosomal marker proteins were higher for the MSC-EV 41.5c preparation than for the MSC-EV 16.3 preparation (Figure 6B). Notably, hardly any differences in the band intensities for MSC-EV 41.5a and b preparations were found. Although the MSC-EV 41.5b preparation contained more particles with a higher purity index than the MSC-EV 41.5a preparation (Figure 6A), MSC-EV 41.5a preparation was identified as being capable of modulating GvHD symptoms at the applied doses, while MSC-EV 41.5b preparation lacked this capability. Thus, none of the applied markers allows any conclusion about the MSC-EV preparations’ therapeutic potential. Consequently, functional analyses remain mandatory to assess the potency of each individual MSC-EV preparation.

**Figure 6:**
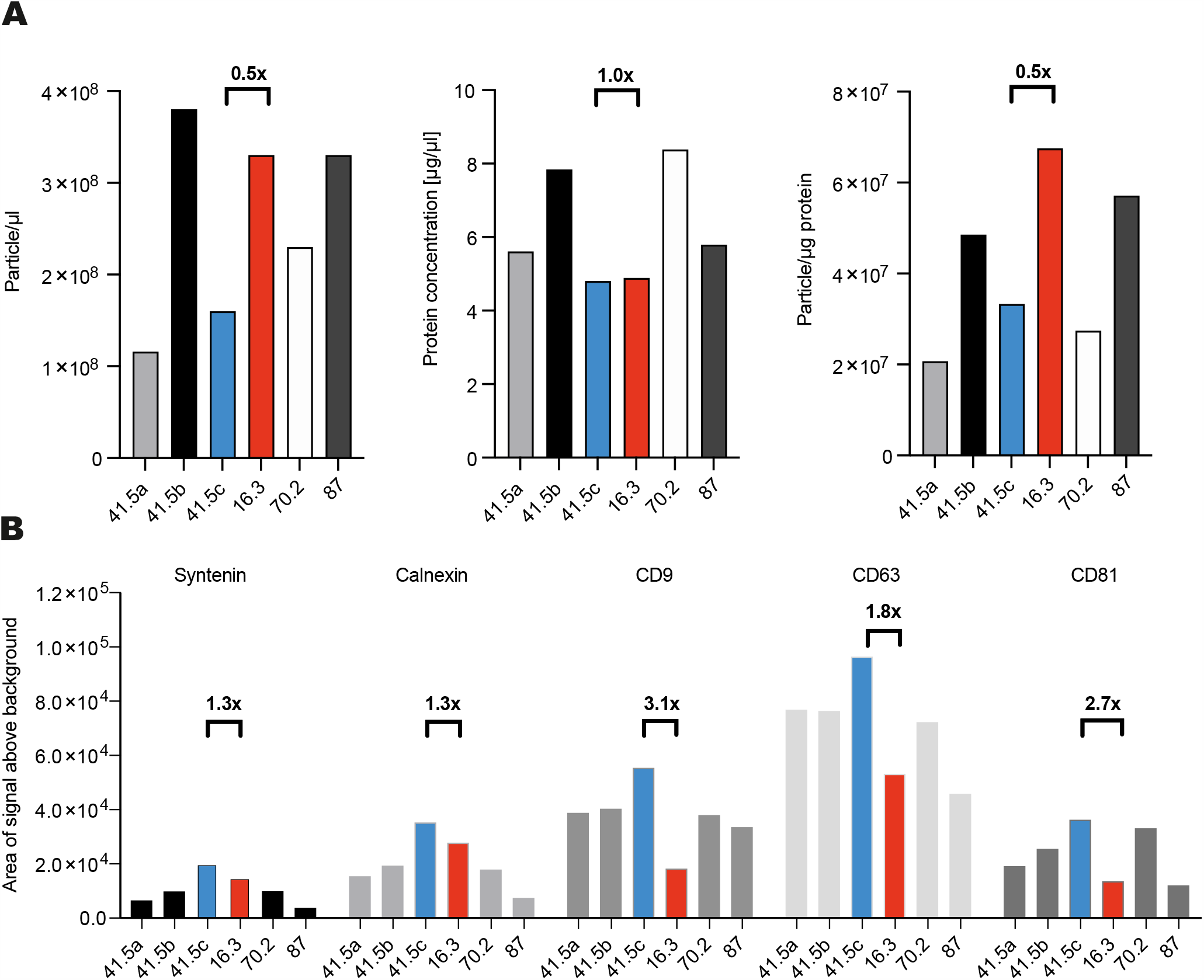
Recorded molecular and physical features of different MSC-EV preparations do not correlate with their functional properties: (A) Particle and protein concentration of the applied MSC-EV preparations were compared; resulting purity indices were calculated as particles per protein amount. (B) Intensities of specific Western blot bands for selected proteins were evaluated semi-quantitatively applying ImageJ analyses on documented Western blot images (suppl. Fig. 2, suppl. Tabel 2). blue, MSC-EV 41.5c preparation as a confirmed active example; red, MSC-EV 16.3 preparation as a confirmed ineffective example.

## Discussion

Inspired by controversies of reported efficacies of MSC therapies especially in steroid refractory aGvHD treatment (Baron and Storb, 2012; Galipeau, 2013; Le Blanc et al., 2004), and given our previous observation that MSC-EV preparations can differ in their cytokine contents (Kordelas et al., 2014), we aimed to comparatively analyse the therapeutic effects of independent MSC-EV preparations in a pre-clinical aGvHD mouse model. To this end, we have adapted and optimized an aGvHD mouse model. Upon co-transplantation of T cell depleted bone marrow cells and defined numbers of purified naive splenic CD4^+^ T cells, very reproducible disease symptoms could be induced. This strategy further improves existing aGvHD mouse models. Until now, for GvHD introduction, either complete bone marrow together with concrete splenic T cell numbers, or T cell depleted BM together with unfractioned splenic cells being adjusted to concrete T cell numbers have been transplanted into myeloablated mice to induce GvHD, respectively (Anderson et al., 2003; Gao et al., 2014; Riesner et al., 2016). In both settings, either due to the presence of BM resident T cells or to residual splenic cells resulting GvHD mice showed more variable disease phenotypes as compared to the advanced setting in our study.

Applying a standardized PEG precipitation protocol involving a final ultracentrifugation step (Borger et al., 2020; Kordelas et al., 2014; Ludwig et al., 2018), EVs were either prepared from supernatants harvested from cultures of different donor derived MSCs or of independent propagations of the same MSC batches. All MSCs fulfilled the *bona fide* ISCT criteria (Dominici et al., 2006). Specifically, all MSC-EV preparations displayed similar particle numbers and expression levels of exosomal marker proteins as determined by immuno blotting. Despite these similarities, merely a proportion of these preparations were found to be therapeutically active. Notably, even EV preparations obtained from conditioned media of independent propagations of the same MSCs variably impacted the aGvHD pathology. Despite the fact that all MSC-EV preparations were produced with the same standardized procedure, our data thus demonstrate functional heterogeneity among independent MSC-EV preparations.

Potentially MSCs might fall into different categories, some of which have capabilities to secrete EVs with therapeutic activities while others fail to secrete such EVs. Apparently, the quality of the MSCs can vary among different donors. In addition, intra-donor variations in MSCs might exist. Upon raising MSCs of individual donors, commonly several independent MSC colonies are formed in the wells original BM specimens have been seeded to. Upon comparing the morphology of MSCs in different colonies, differences in their sizes as well as in their migration, adhesion and proliferation capabilities can be observed. Thus, MSCs raised from primary material regularly provide an oligo-clonal mixture of different MSC subtypes. Hypothetically, some propagated MSC cultures may contain a higher proportion of MSC subtypes that secrete therapeutically active EVs than others. The stoichiometry among such different subtypes likely is stochastic within the MSC culture and may vary from one to the other MSC propagation. For now, we are not aware of any surrogate marker allowing the discrimination of functionally different MSC subtypes towards evaluation of such putative stoichiometry in MSC products.

Conceding an issue of MSC heterogeneity, which can also severely be influenced by MSC growth conditions (Reiner et al., 2017), it is also non-deducible from the MSCs phenotypes, yet, whether resulting EV preparations have a higher or lower chance to be therapeutically active. Also at the EV level we have not identified any surrogate parameters, which would allow us to discriminate more potent from less potent MSC-EV preparations. Thus, to avoid any failures in proposed clinical MSC-EV trials, we see it as mandatory that each MSC-EV preparation that has been produced is tested for its potency in an appropriate assay. To this end, we need to admit that potency testing of given MSC-EV preparations is challenging itself. We have recently reviewed such challenges in detail (Gimona et al., Cytotherapy in revision). Briefly, the mechanism of action (MoA) of MSC-EVs needs to be considered to be multimodal; in addition to immunomodulatory properties, MSC-EV preparations have been reported to exert pro-angiogenic, pro-regenerative and anti-apoptotic activities (Arslan et al., 2013; Börger et al., 2017). It might depend on the target disease which of those activities might be more important to successfully suppress disease symptoms. Knowledge about the MoA most certainly will help to identify critical activities or even surrogates being essential to mediate such activities. However, for now, we have not identified the side of action (SoA) of MSC-EV preparations, further challenging the dissection of the MoA with all its modalities. To this end, it is often assumed that applied MSC-EV preparations act in lesion sites or disease affected tissues, respectively. However, this hypothesis may not be universally true. Exemplified by an ischemic stroke model, we have demonstrated that MSC-EVs, which successfully suppress ischemic stroke symptoms, significantly affect the immune system of respective animals (Doeppner et al., 2015; Wang et al., 2020). On the one hand they repress stroke induced lymphopenia (Doeppner et al., 2015) and on the other hand immune cell infiltration, especially of lymphocytes, monocytes and neutrophils into the lesion sites (Wang et al., 2020). The observation that neutrophil depletion and MSC-EV administration mediate comparable, non-synergistic neuroprotective effects (including suppression of leukocyte migration into lesion sites) suggests a central role of neutrophils in disease progression and implies an impact of immunomodulatory EVs on neutrophilsat at least in ischemic stroke (Wang et al., 2020). Thus, we consider that the MSC-EV preparations’ direct SoA is somewhere in the periphery rather than in the brain. The issue is even more complex given no existing EV preparation method really isolates EVs. For now we rather use EV enrichment procedures all resulting in containment of by-products that might importantly contribute to the MoA of obtained EV preparations (Witwer et al., 2019). Accordingly, it has been recently reported that pro-angiogenic activities in MSC-EV samples might be mediated by non-EV associated components (Whittaker et al., 2020). Anyway, whatever the MoA, the SoA or the active components of given MSC-EV preparations are, if containing the right activity they appear as very powerful therapeutics for many diseases including treatment for COVID-19 patients (Börger et al., 2020). To not repeat failures in the MSC field, i.e. the failure of a phase III clinical trial for MSCs in aGvHD treatment (Galipeau, 2013), we do highly recommend to include evaluated assays whose results correlate with outcomes in preclinical models. Here, we have used a novel form of a MLR assay whose prediction for immunomodulatory capabilities of given MSC-EV preparations correlated very well with observed effects in aGvHD mice. Although we consider that the MLR assay reveals essential information about the potency of respective MSC-EV preparations, it is not a potency assay. The term “potency assay” is a regulatory term and potency assays need to fulfil several requirements of regulatory authorities (Gimona et al, Cytotherapy in Revision). It needs to be standardized and evaluated, which according to our understanding would be very challenging for an assay, which is based on primary human cells. Thus, for now, it remains an important challenge in the field to increase our overall understanding of the MSC-EVs therapeutic potentials and to develop appropriate, evaluated potency assays which allow us to discriminate therapeutic active from non-active MSC-EV products in the future. Since we lack a lot of knowledge about our products, it is absolutely required that production and characterisation details of clinically applied MSC-EV preparations are appropriately reported, especially if clinical administration of MSC-EV products is published in peer-reviewed journals. Besides the MISEV2018 and EV-TRACK recommendations (Consortium et al., 2017; Thery et al., 2018), as part of a panel of experts, we strive for identification of critical parameters in the MSC-EV field which should be considered and appropriately reported (Lener et al., 2015; Reiner et al., 2017; Witwer et al., 2019). We guess the results of this study emphasize the substantial therapeutic potential that can be provided by appropriate MSC-EV products. However, our results also demonstrate that MSC-EV products should not be considered as effective across the board and several challenges need to be encountered before translating MSC-EVs into the clinics.

## Acknowledgments

We thank the Westdeutsche SpenderZentrale (WSZE) for providing bone marrow samples of healthy donors. We are also grateful to all healthy blood donors whose cells were used in the MLR assay. This study was supported by the LeitmarktAgentur.NRW and the European Union (European Regional Development Fund 2014-2020, EFRE-0800396), the Stem Cell Network North Rhine Westphalia and the European Union (ERA-NET EuroTransbio 11: EVTrust [031B0332B]; EU COST program ME-HaD [BM1202]). We thank Thomas Scholtysik for technical support in the GvHD experiments.

## Declaration of Interest Statement

BG is a scientific advisory board member of Evox Therapeutics and Innovex Therapeutics SL. All other authors report no conflicts of interest.

## Appendices

## Supplement figures

**Suppl.-Figure 1:**
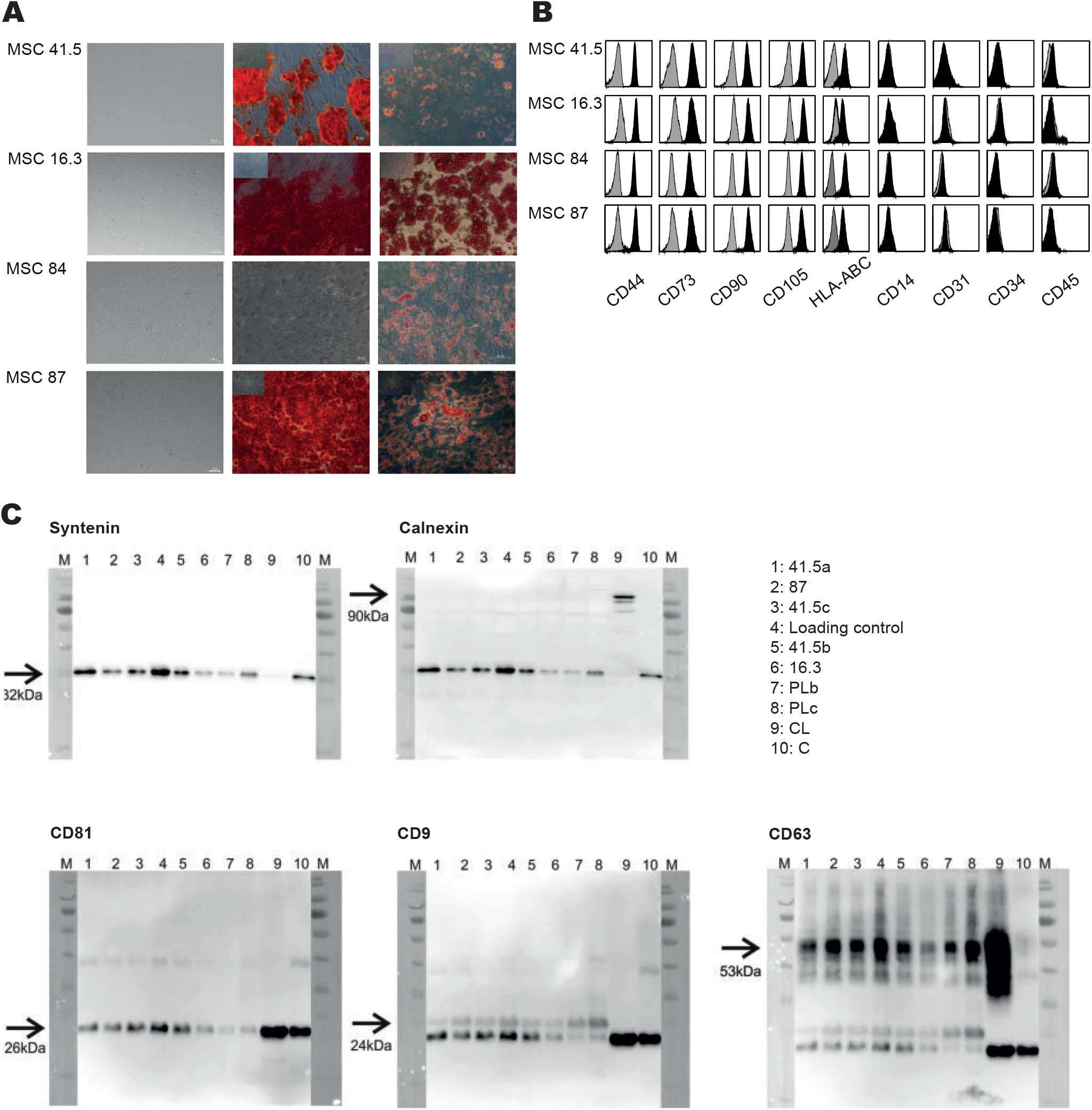
All MSCs provide MSC *bona fide* criteria. (A) Morphological appearance of expanded MSCs (phase contrast image, first column) and their osteogenic (2^nd^ column) and adipogenic differentiation potential (3^rd^ column) following alizarin red or oil red staining, respectively. Inserts in the 2^nd^ and 3^rd^ column show negative controls. (B) cell surface phenotype of expanded MSCs analysed by flow cytometry. Cells were labelled with fluorochrome conjugated antibodies against the MSC marker proteins CD44, CD73, CD90, CD105 and HLA-ABC and the negative markers CD14, CD31, CD34 and CD45.

**Suppl.-Figure 2:** All MSC-EV preparations contain exosomal marker proteins and lack the impurity marker Calnexin. (A) Western blots of all MSC-EV preparations used in the study. The plot in the upper row was initially stained with anti-Synthenin antibodies. Following documentation the blot was additionally labelled with anti-Calnexin antibodies (no stripping). The plot in the lower row was sequentially stained with anti-CD81, anti-CD9 and anti-CD63 antibodies. Results after each detection round are depicted. Lane 9 contains MSC lysates (CL) and lane 10 a non-MSC derived EV preparation we internally use as a CD81 positive control.

## Supplemental Tables

**Suppl. Table 1:**
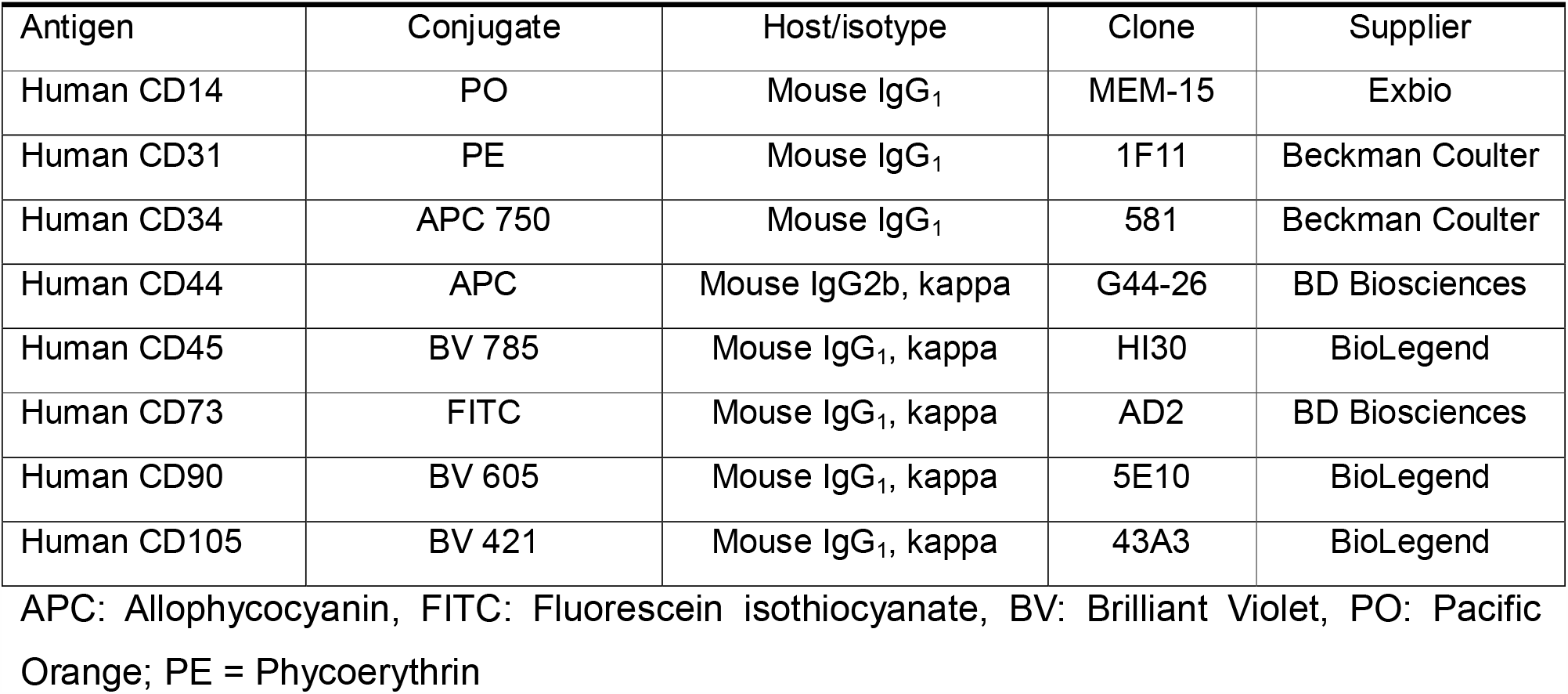
Applied fluorescence conjugated antibodies

**Suppl.-Table 2:**
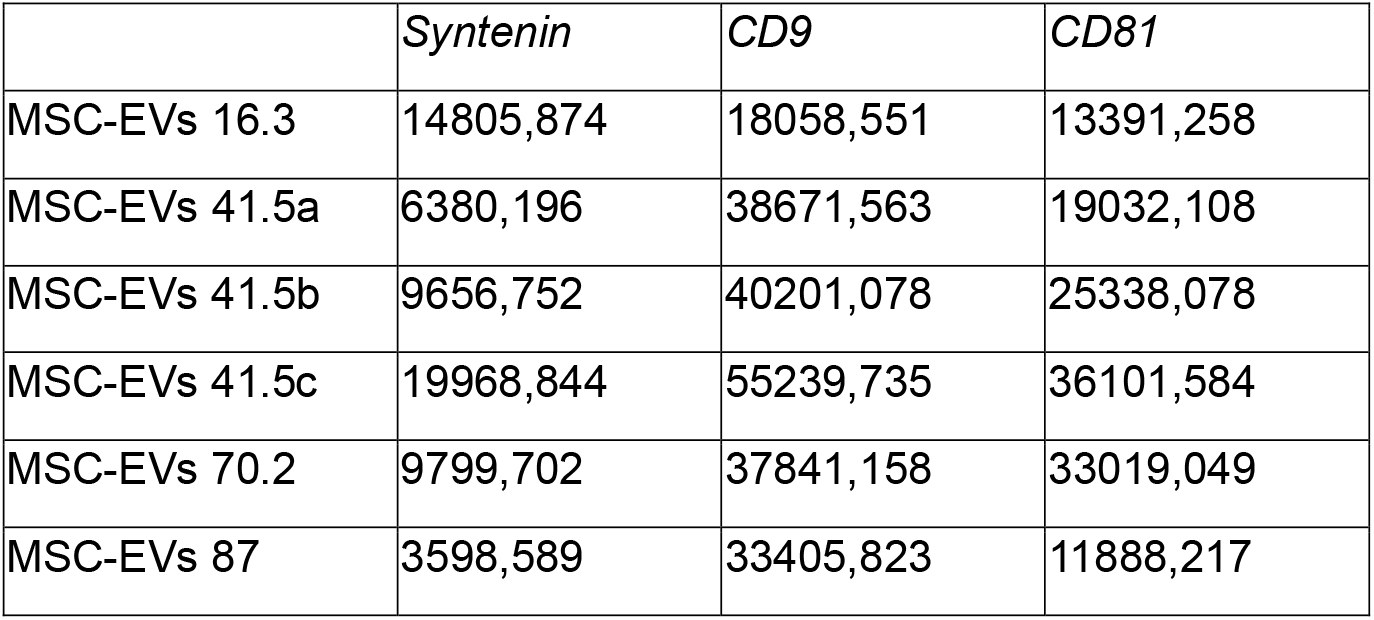
Band intensities of WB shown in Suppl. Fig. 2 as quantified by Image J.

