## Supplemental figures for "Independent human mesenchymal stromal cell-derived extracellular vesicle preparations differentially affect symptoms in an advanced murine Graft-versus-Host-Disease model"

### Supplemental Tables:

**Suppl.-Table 1:** Applied fluorescence conjugated antibodies

| Antigen | Conjugate | Host/isotype | Clone | Supplier |
|---|---|---|---|---|
| Human CD14 | PO | Mouse $IgG_1$ | MEM-15 | Exbio |
| Human CD31 | PE | Mouse $IgG_1$ | 1F11 | Beckman Coulter |
| Human CD34 | APC 750 | Mouse $IgG_1$ | 581 | Beckman Coulter |
| Human CD44 | APC | Mouse IgG2b, kappa | G44-26 | BD Biosciences |
| Human CD45 | BV 785 | Mouse $IgG_1$, kappa | HI30 | BioLegend |
| Human CD73 | FITC | Mouse $IgG_1$, kappa | AD2 | BD Biosciences |
| Human CD90 | BV 605 | Mouse $IgG_1$, kappa | 5E10 | BioLegend |
| Human CD105 | BV 421 | Mouse $IgG_1$, kappa | 43A3 | BioLegend |

APC: Allophycocyanin, FITC: Fluorescein isothiocyanate, BV: Brilliant Violet, PO: Pacific Orange; PE = Phycoerythrin

**Suppl.-Table 2:** Band intensities of WB shown in Suppl. Fig. 2 as quantified by Image J.

| | *Syntenin* | *CD9* | *CD81* |
|---|---|---|---|
| MSC-EVs 16.3 | 14805,874 | 18058,551 | 13391,258 |
| MSC-EVs 41.5a | 6380,196 | 38671,563 | 19032,108 |
| MSC-EVs 41.5b | 9656,752 | 40201,078 | 25338,078 |
| MSC-EVs 41.5c | 19968,844 | 55239,735 | 36101,584 |
| MSC-EVs 70.2 | 9799,702 | 37841,158 | 33019,049 |
| MSC-EVs 87 | 3598,589 | 33405,823 | 11888,217 |

### Supplement figures

**Suppl.-Fig. 1**

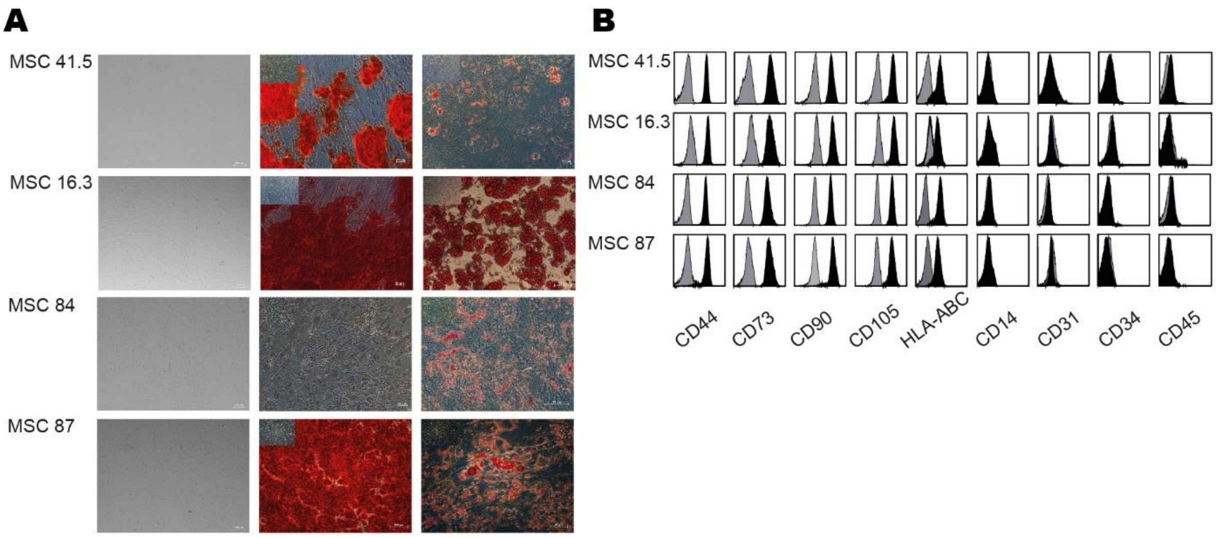


**Suppl.-Figure 1: All MSCs provide MSC *bona fide* criteria.** (A) Morphological appearance of expanded MSCs (phase contrast image, first column) and their osteogenic (2nd column) and adipogenic differentiation potential (3rd column) following alizarin red or oil red staining, respectively. Inserts in the 2nd and 3rd column show negative controls. (B) cell surface phenotype of expanded MSCs analysed by flow cytometry. Cells were labelled with fluorochrome conjugated antibodies against the MSC marker proteins CD44, CD73, CD90, CD105 and HLA-ABC and the negative markers CD14, CD31, CD34 and CD45.

**Suppl.-Fig. 2**

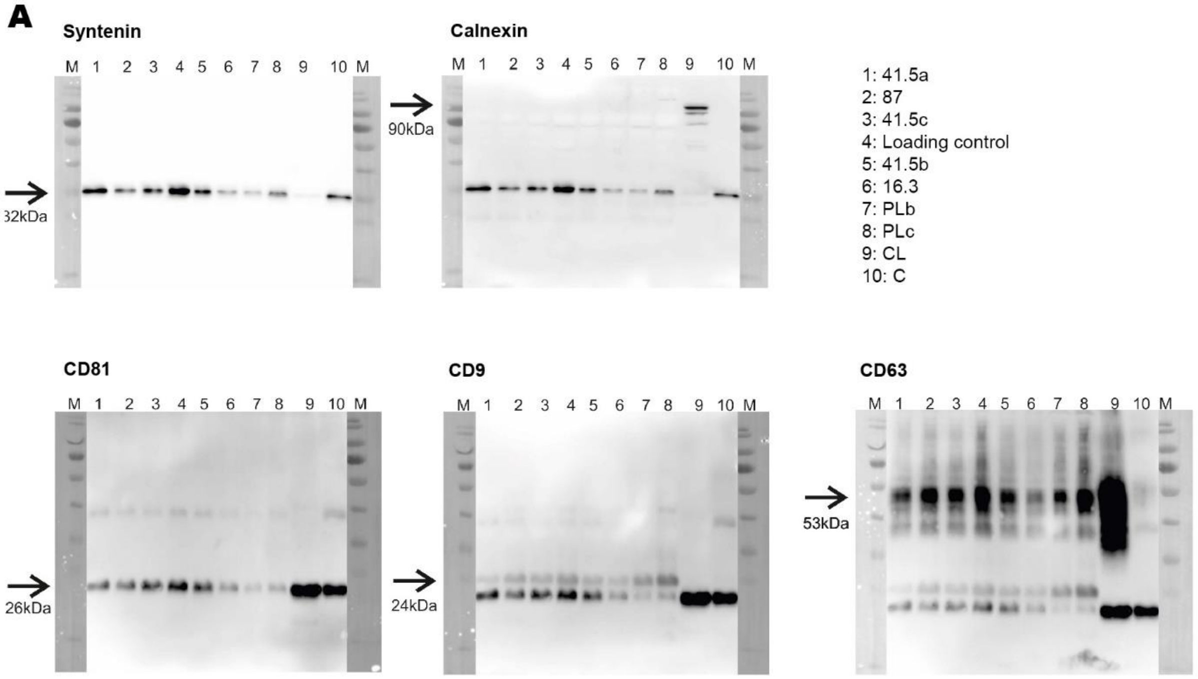


**Suppl.-Figure 2: All MSC-EV preparations contain exosomal marker proteins and lack the impurity marker Calnexin.** (A**)** Western blots of all MSC-EV preparations used in the study. The plot in the upper row was initially stained with anti-Synthenin antibodies. Following documentation the blot was additionally labelled with anti-Calnexin antibodies (no stripping). The plot in the lower row was sequentially stained with anti-CD81, anti-CD9 and anti-CD63 antibodies. Results after each detection round are depicted. Lane 9 contains MSC lysates (CL) and lane 10 a non-MSC derived EV preparation we internally use as a CD81 positive control.

<u>**Supplement figures**</u>

**Suppl.-Figure 1: All MSCs provide MSC *bona fide* criteria.**

**Suppl.-Figure 2: All MSC-EV preparations contain exosomal marker proteins and lack the impurity marker Calnexin.**
